# Performance Comparison Of Agilent New SureSelect All Exon v8 Probes With v7 Probes For Exome Sequencing

**DOI:** 10.1101/2022.04.15.488338

**Authors:** Vera Belova, Anna Shmitko, Anna Pavlova, Robert Afasizhev, Valery Cheranev, Anastasia Tabanakova, Natalya Ponikarovskaya, Denis Rebrikov, Dmitriy Korostin

## Abstract

Exome sequencing may become routine in health care it increases the chance of pinpointing the genetic cause of an individual patient’s condition and thus making an accurate diagnosis. It is important for facilities providing genetic services to keep track of changes in the technology of exome capture in order to maximize throughput while reducing cost per sample. In this study, we focused on comparing the newly released exome probe set Agilent SureSelect Human All Exon v8 and the previous probe set v7. In preparation for higher throughput of exome sequencing using the DNBSEQ-G400, we evaluated target design, coverage statistics, and variants across these two different exome capture products. Although the target size of the v8 design has not changed much compared to the v7 design (35.24 Mb vs 35.8 Mb), the v8 probe design allows you to call more of SNVs (+3.06%) and indels (+8.49%) with the same number of raw reads per sample on the common target regions (34.84 Mb). Our results suggest that the new Agilent v8 probe set for exome sequencing yields better data quality than the current Agilent v7 set.

## Introduction

Whole exome sequencing (WES) is widely used in genomic studies as well as genetic tests. Exons (protein coding regions) represent 1-2% of the human genome comprising up to 85% of the known variants significant for diagnostics (1). At the same time, WES is 3-5 times cheaper than whole genome sequencing (2). Currently, exome analysis, embracing a set of different characteristics, has proven to be a more efficient diagnostic tool being especially effective in the area of human clinical genetics (3).

There are several commercial kits for whole exome enrichment. The most known kits are SureSelect (Agilent), TruSeq Capture (Illumina), xGen (IDT), Human Comprehensive Exome (Twist Bioscience), SeqCap EZ (Roche NimbleGen) (4-9). Enrichment protocols are similar and are based on hybridization of exon sequences with biotinylated DNA or RNA probes with a subsequent capture by streptavidin-covered magnetic beads. Most kits are designed to enrich the libraries for sequencing using the Illumina platform. However, earlier, we managed to adapt the enrichment protocol for sequencing using the MGI platform with the Agilent SureSelect Human All Exon V6 probes.

In 2021, Agilent launched an updated enrichment probe set v8 and compared its performance with the other manufacturer (10) but not with the previous version v7. In this study, we focused on comparing the Agilent SureSelect Human All Exon v7 and v8 designed probes. We studied the changes introduced in panel design, metrics of enrichment quality and statistically assessed the efficiency and quality of the variant detection (11).

We prepared 20 libraries, divided them into 2 pools of 10 libraries and performed 2 rounds of enrichment of the pools using the v7 or v8 probes following the RSMU_exome protocol (12). The sequenced pools were compared using bioinformatics pipeline based on the following characteristics: target regions, the percentages of on-targets, off-targets, and duplicates as well depth of coverage of the regions with various GC content.

## Materials and methods

### Ethics Statement

This study conformed to the principles of the Declaration of Helsinki. The appropriate institutional review board approval for this study was obtained from the Ethics Committee at the Pirogov Medical University. All patients provided informed consent for sample collection, subsequent analysis, and publication thereof.

### Sample Preparation and Sequencing

The libraries were prepared from 20 samples containing 300-600 ng of human genomic DNA taken from 20 patients using MGIEasy Universal DNA Library Prep Set (MGI Tech) following the manufacturer’s instructions. DNA fragmentation was performed by sonication with the average fragment length of 250 bp using Covaris S-220. Quality control of the obtained DNA libraries was performed using the High Sensitivity DNA assay with the 2100 Bioanalyzer System (Agilent Technologies).

Previously pooled DNA libraries were enriched following the RSMU_exome protocol (12). 20 DNA libraries were divided into 2 pools each containing 10 libraries. Each pool was enriched twice with the SureSelect Human All Exon v7 probes and the latest version of the probes SureSelect Human All Exon v8 (Agilent Technologies) for the second time. Finally, we obtained 4 enriched DNA library pools. The concentrations of the prepared libraries were measured using Qubit Flex (Life Technologies) with the dsDNA HS Assay Kit. The quality of the prepared libraries was assessed using Bioanalyzer 2100 with the High Sensitivity DNA kit (Agilent Technologies).

The enriched library pools were further circularised and sequenced by a paired end sequencing using DNBSEQ-G400 with the High-throughput Sequencing Set PE100 following the manufacturer’s instructions (MGI Tech) with the average coverage of 100x. We loaded one pool per lane into the patterned flow cells in two different runs. FastQ files were generated using the zebracallV2 software by the manufacturer (MGI Tech).

### Bioinformatics pipeline

The quality of the obtained 40 paired fastq files was analysed using FastQC v0.11.9 (13). Based on the quality metrics, the fastq files were trimmed using Trimmomatic v0.39 (14). To correctly estimate the enrichment and sequencing quality, all 20 exomes were downsampled to 50 million reads using Picard DownsampleSam v2.22.4 (15). Reads were aligned to the indexed reference genome GRCh37 using bwa-mem (16). SAM files were converted into BAM files and sorted using SAMtools v1.9 to check the percentage of the aligned reads (17). Based on the obtained BAM files, the quality metrics of exome enrichment and sequencing were calculated using Picard v2.22.4, and the number of duplicates was calculated using Picard MarkDuplicates v2.22.4. We performed the quality control analysis with the following bed files: Agilent v7_regions, Agilent v8_regions. Bed files for the GENCODE and RefSeq databases were uploaded from the UCSC Table Browser (https://genome.ucsc.edu/cgi-bin/hgTables?hgsid=1309831311_Di0qVAk2HAMSBFgug0SoMWuDiYQT). Genomic coordinates of unique v7 and v8 regions in the bed files were annotated using the Panther database (18). Variant calling was performed using bcftools mpileup v1.9.

## Results

### Comparison of probe designs

We detected several changes in target design of v8 compared to v7 introduced by the manufacturer. The manufacturer did not alter the probe structure preserving 120 bp biotinylated cRNA probes. The manufacturer claims that coding content was updated according to the database releases (CCDS release 22, GENCODE V31, RefSeq release 95), added the TERT promoter region, but removed non-coding ClinVar Pathogenic variants. The target size of the v8 kit is 35.24 Mb, whereas the target size of the v7 kit is 35.8 Mb, the intersection of the bed files from both kits is 98.42% (34.84 Mb). The percentage of unique target regions is 2.69% (0.96 Mb) and 1.14% (0.4 Mb) for the Agilent v7 and v8 exome, respectively. We compared the v7 and v8 bed files with the bed file containing the coding exons of the GENCODE Genes track (basic subtrack, release V39lift37, Oct 2021) (34.93 Mb). The intersection between v8 and GENCODE v39 was 98.9% (34.07 Mb), the intersection between v7 and GENCODE v39 was 98.2% (34.3 Mb), and 0.29 Mb of the GENCODE v39 regions were absent in both kits.

We collected precise information (chromosomic coordinates, Gene ID, an annotation) on target regions from the v7 (0.96 Mb) and v8 kits (0.4 Mb) which is provided in the Supplementary Table 1. The plot analysing the distribution of lengths of the changed fragments (Supplementary Table 1) demonstrates that most altered positions are short (less than several dozens of base pairs) which means that the manufacturer adjusted design of certain probes using the previous version of the targets. We analysed those fragments that were longer than **30 bp** as we were interested in detecting unique fragments for the v7 and v8 kits in the current version of the bed file of the GENCODE v39 database. Our analysis also included the bed file containing the coordinates of all exons plus 20 bases at each end from the RefSeq ALL database (Source data version: NCBI Homo sapiens 109.20211119 (2021-11-23)). The percentage of the included bed with unique fragments of the v8 target into the current GENCODE v39 bed file was higher than that of the previous version v7 as expected. The intersection of the unique regions of the current GENCODE v39 with v8 was 0.33 Mb and with v7 was 0.09 Mb. The intersection of the RefSeq bed with a larger size (95.47 Mb) which includes exons +-20 bp from all curated and predicted genes with v8 was 0.23 Mb and with v7 0.26 Mb.

### Enrichment quality

To assess the enrichment quality, the obtained data (raw reads) for 40 exomes (20 samples enriched by the v7 or v8 probes) were downsampled to 50M reads. The coverage statistics was calculated using Picard, and metrics were averaged for the samples from each v7 and v8 pool. The results obtained for each downsampled sample in the pool are shown in the Supplementary Table 2. We detected no significant differences in the number of on-target and off-target reads and the percentage of duplicates. The average values are within the range of 0.2-1.3% for both v7 and v8 (Fig.2). Mean target coverage was similar for both kits and was 56.38x and 56.88x for v7 and v8, respectively. However, median target coverage values for two kits were different and equal to 53.4x and 48.6x for v8 and v7, respectively. Therefore, the dispersion of target coverage values for v8 is lower indicating a higher coverage uniformity.

**Figure 1.**
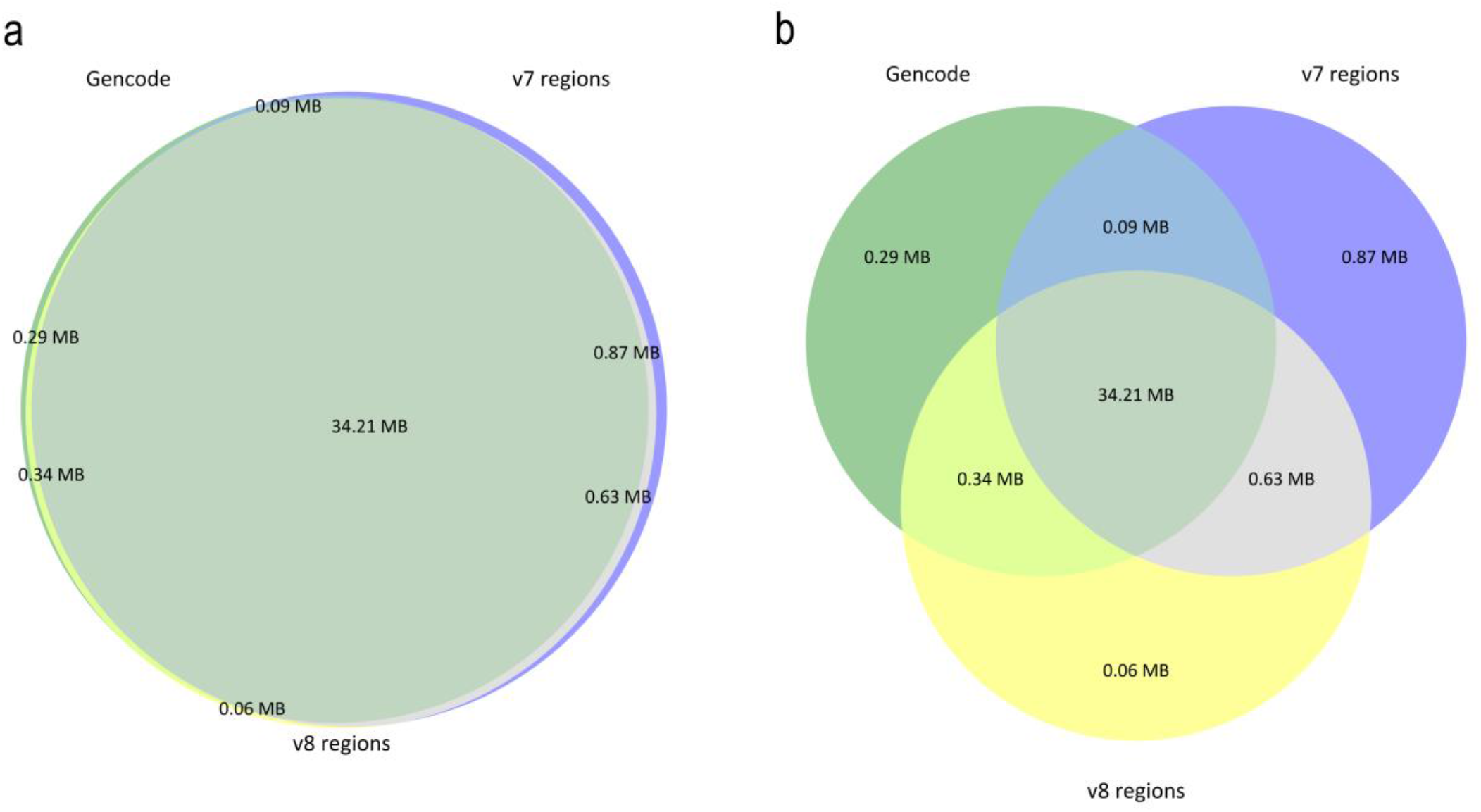
Venn diagram showing the intersection between the Agilent v7 exome (35.8 Mb), Agilent v8 exome (35.24 Mb), and Gencode V39 coding exons (34.93 Mb): A) weighted, B) unweighted.

**Figure 2.**
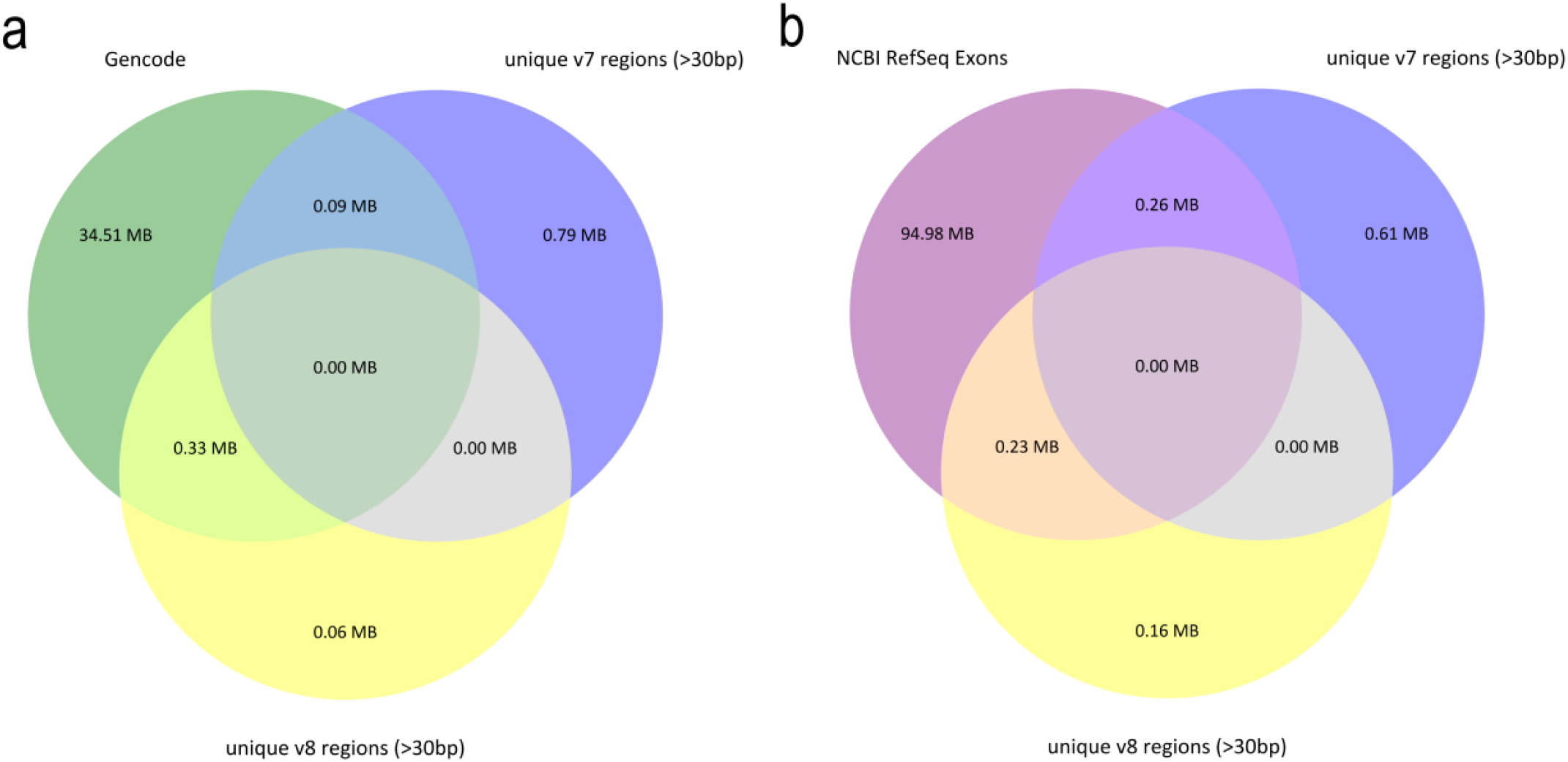
Venn diagram showing the intersection of unique regions of the Agilent v7 exome (0.96 Mb) and Agilent v8 exome (0.4 Mb) with: A) Gencode V39 coding exons (34.93 Mb); (B) NCBI RefSeq ALL exons +-20bp (95.47 Mb).

**Figure 3.**
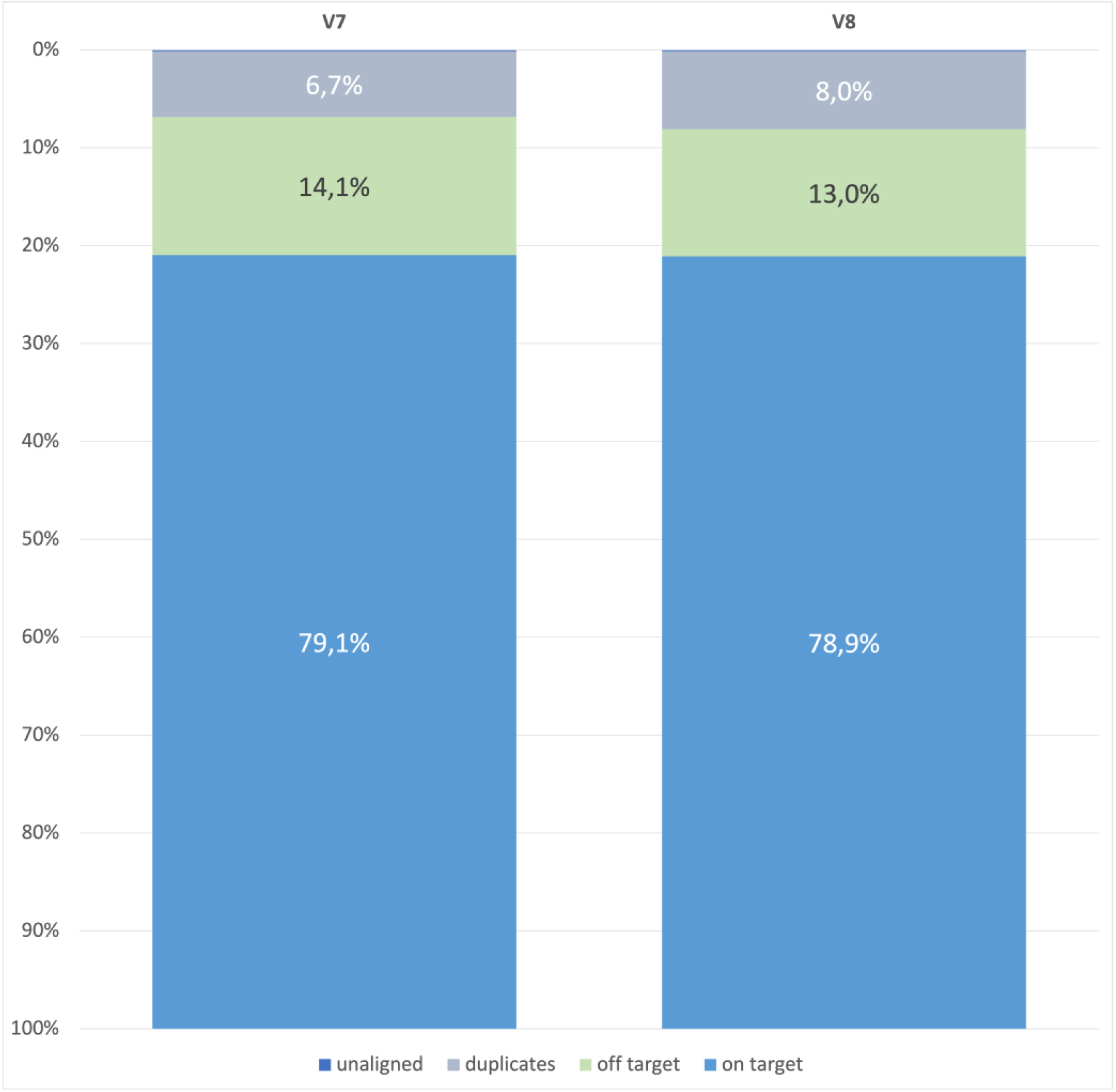
Stacked barplots show the average values for on-targets, off-targets, duplicates, and unaligned reads in the samples from the v7 and v8 exome pools (downsampled results).

The average values of metrics in the pools which reflect the target region coverage quality in the v8 kit are higher than in the v7 kit. The percentage of the target regions with ≥10x coverage is 96.28% and 95.08% for v8 and v7, respectively. At the same time, the percentage of the target regions with 20x coverage in v7 is 5% less than in v8 (88.07% for v7 vs. 92.96% for v8) indicating higher enrichment quality obtained with the v8 probes. The 40x on-target coverage is in the range of 57-66% (mean = 62%) for the v7 kit and is 10% less than that of the v8 kit (which is in the range of 65%-77%, mean = 71%). At the same time, the distribution of v8 is closer to the normal distribution, the position coverage is more uniform, there are fewer overcovered (≥80x coverage) or undercovered positions (the inflection point is shown in Fig.4A) which allows obtaining the sufficient coverage for more positions using less data.

**Figure 4.**
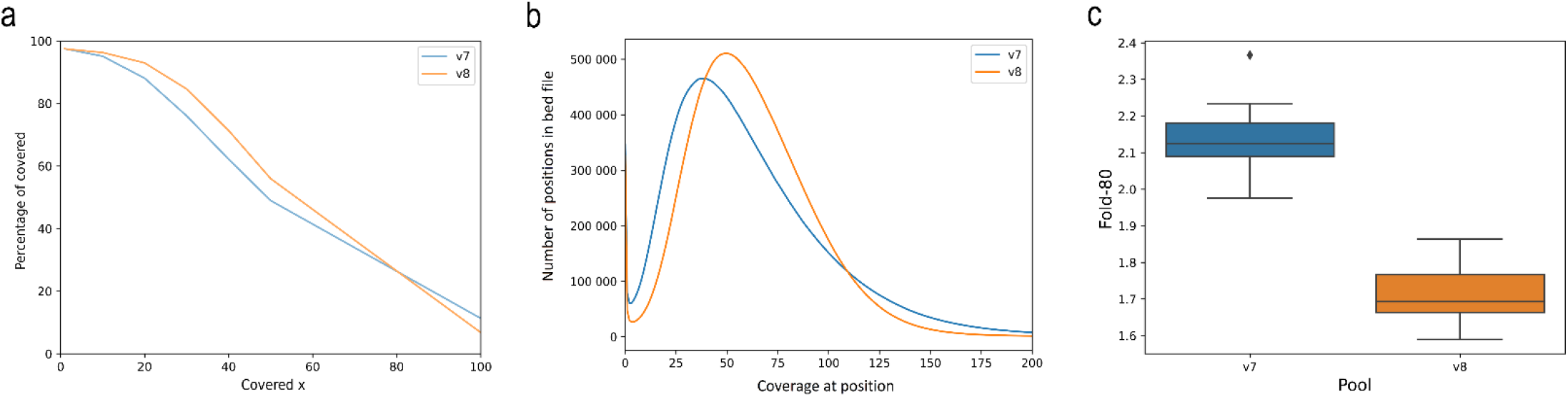
(A,B,C). Performance of exome protocol in terms of coverage quality in downsampled samples: A) Dependence of coverage quality of the v7 and v8 target regions from depth; B) Graph showing the number of positions (bp, y-axis) vs. the coverage (x-axis); C) Box plot showing the Fold-80 metric for the samples from the v7 and v8 pools.

The FOLD_80 parameter which reflects the coverage uniformity in the v8 pool samples (mean = 1.72) is better than that of the v7 pool samples (mean = 2,13) (Fig.4C). The closer the value is to 1, the fewer rounds of sequencing a sample requires to obtain 80% of the targeted bases with the original mean coverage.

### GC content

The AT_DROPOUT metric is 2 times lower for the exomes enriched with the v8 kit (v7 mean = 29.23%, v8 mean = 15.92%). GC_DROPOUT does not differ between the kits (v7 mean = 12.9%, v8 mean = 13.09). Both AT_DROPOUT and GC_DROPOUT metrics indicate the percentage of misaligned reads that correlate with low (%-GC is < 50%) or high (%-GC is > 50%) GC content, respectively. Fig.5 demonstrates that the v8 probes provide slightly more uniform coverage of regions with the GC content in the range of 40-60% (the red zone in the figures). However, this value is high in the v7 probes as well.

**Figure 5.**
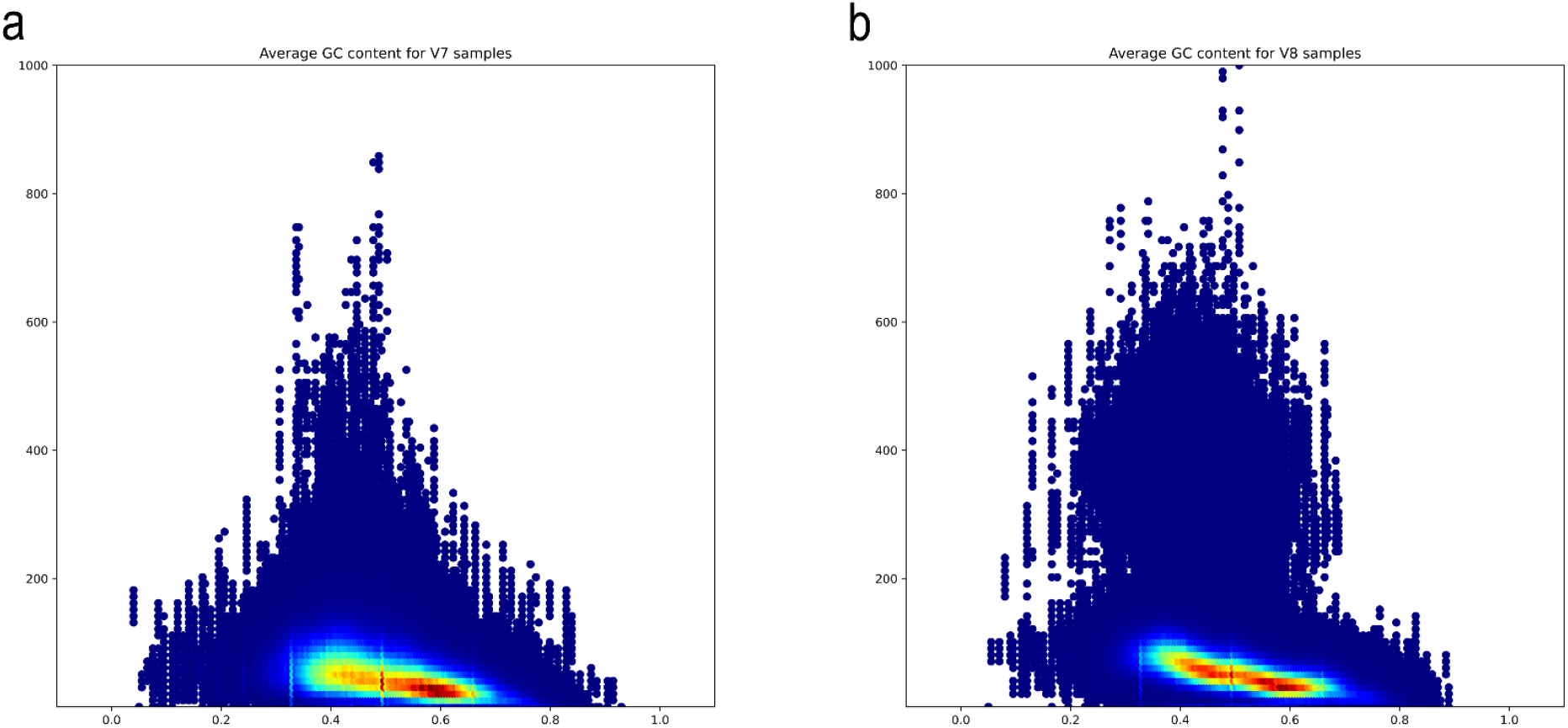
Density plot showing %GC content vs. Mean Depth for: (A) Agilent v7; (B) Agilent v8. Density plot showing %GC content vs. mean depth. The data in this plot was collected by merging all samples from the V7 and V8 pools. Density estimation was performed using 2D plots. More specifically, we chose data points in fixed rectangle (%GC content ∈[0;1], mean depth ∈[0;1000]) and split it into evenly spaced 200×100 grid and counted the data points in each cell of the grid. Finally, we normalised the grid to the range of [0,1] and plotted it using “jet” colormap from matplotlib library.

### SNV and INDEL calling comparison

Furthermore, we estimated the calling quality by calculating the number of single nucleotide variants (SNVs) and small insertions and deletions (indels) detected by different kits with the equal number of raw reads per sample. Table 1 shows the average result of calling for the v7 and v8 pools filtered by the quality of entire bed files (results for each sample are provided in Supplementary Table 3). The following filters were used for calling: cut-off for the variants with the coverage depth exceeding 13 reads (DP > 13) and a parameter QUAL > 30. The average numbers of SNVs and indels obtained were 25’736 and 743 for the v7 exomes and 25’558 and 699 for v8. A higher amount of called variants for the v7 kit can be accounted for a larger size of target design. As different kits provide bed files with different sizes, we compared variant calling in the overlapping target regions of the v7 and v8 kits. This approach enables correct comparison of two probe designs. Using the same target (bed V7 cross v8 = 34.84 Mb), we calculated the average variant numbers. The number of SNVs and indels for the samples from the v8 pool were 3.06% and 8.49% higher than that of the v7 pool, respectively (Table 1).

**Table 1.** Average results of variant calling of SNV and indels for the samples from the v7 and v8 pools using their own target (bed v7, bed v8) and target intersection (bed v7 vs. v8) filtered by DP > 13 and QUAL>30.

| Design | Variant type | Count on target v7<br>(35.7 Mb) | Count on target v8<br>(35.1 Mb) | Count on target<br>V7 cross v8<br>(34.84 Mb) |
| --- | --- | --- | --- | --- |
| V7 pool | SNV | 25 736 | - | 24 374 |
|  | indel | 743 | - | 612 |
| V8 pool | SNV | - | 25 558 | 25 120 **(+3.06%)** |
|  | indel | - | 699 | 664 **(+8.49%)** |

## Discussion

Overall, 2.76% of the target was excluded from v7, while 1.15% of the target was included into v8. Based on our data, we believe that no dramatic changes in probes significant for the clinical potential were added. Most modifications probably lie in changing the approach powered by machine learning to probe design. Most changed fragments in certain regions are several base pair long thus implying that manufacturer inly adjusted certain target regions. However, some targets were quite long (dozens to thousands of base pairs) which indicates the functional changes as well. Changes affecting longer fragments arise from the updated information in the current versions of the databases. For instance, the transcript of the largest fragment in the NACA gene (3330 bp) that was excluded from the v8 target undergoes splicing and is characterised as tsl5 – no single transcript supports the model structure (ENST00000454682.6 NACA-203). The manufacturer excluded all regions now considered to be not protein coding from the target and included certain regions that were unknown when the v7 probes were designed. This can be proved by analysing the intersection between the unique regions of both kits and the latest releases of databases in the same way we analysed Gencode V39 release (Oct 2021).

The major problem of WES is a non-uniform coverage of target regions resulting from the sensitive hybridization reaction of probes with the target fragments of DNA libraries. The introduced changes in the v8 probe design markedly improved the enrichment quality. The v8 probes with the same sizes of raw data per sample provided higher coverage of the larger percent of target regions. We noted that the degree of inadequate coverage of a particular region based on its AT content was better in case of the v8 version. We wondered if variant calling detected the same SNVs and indels in the samples obtained with the v7 and v8 kits. Indeed, the samples enriched with the v8 probes allowed for obtaining more useful data than the v7 probes due to new probe design and higher enrichment quality (uniform coverage of target fragments).

Noteworthy, the presented calculations were performed according to our in-lab gDNA standards. We aimed at estimating relative statistical metrics rather than absolute metrics as it is more correct to analyse them similarly to GIAB or Platinum Genomes. We intended to reveal the advantages that could be gained by an NGS facility performing exome sequencing if it switched to a new version of an enrichment kit.

Therefore, novel probe design Agilent all-exon v8 provides enough advantages as compared to the previous version of the kit and can be recommended as an advanced, more efficient generation of sequencing kits.

## Data availability

All 40 exome sequences will be deposited into the NCBI open-access sequence read archive (SRA) in fastq.gz format. BioProject number will be added as soon as possible.

## Author contributions

VB – Conceptualization, Methodology, Investigation, Validation, Writing – Original Draft Preparation; ASh – Methodology, Investigation, Writing – Original Draft Preparation; AP and RA – Formal Analysis, Methodology, Software, Visualization; VCh, AT, NP – Investigation; DR – Resources and Funding Acquisition; DK – Conceptualization, Project Administration, Methodology, Supervision, Writing – Review & Editing.

